# Genome-wide association study of delay discounting in Heterogenous Stock rats

**DOI:** 10.1101/2023.12.12.570851

**Authors:** Montana Kay Lara, Apurva S. Chitre, Denghui Chen, Benjamin B. Johnson, Khai-Minh Nguyen, Katarina A. Cohen, Sakina A. Muckadam, Bonnie Lin, Shae Ziegler, Angela Beeson, Thiago Sanches, Leah C. Solberg Woods, Oksana Polesskaya, Abraham A. Palmer, Suzanne H. Mitchell

**Affiliations:** Department of Psychiatry, University of California San Diego, La Jolla, CA, 92093, USA; Department of Internal Medicine, Wake Forest School of Medicine, Winston-Salem, NC, 27157, USA; Institute for Genomic Medicine, University of California San Diego, La Jolla, CA, 92093, USA; Departments of Behavioral Neuroscience, Psychiatry, the Oregon Institute of Occupational Health Sciences, Oregon Health & Science University, Portland, OR, 97239 USA

**Keywords:** Delay discounting, adjusting amount, GWAS, Heterogenous Stock rats

## Abstract

Delay discounting refers to the behavioral tendency to devalue rewards as a function of their delay in receipt. Heightened delay discounting has been associated with substance use disorders, as well as multiple co-occurring psychopathologies. Genetic studies in humans and animal models have established that delay discounting is a heritable trait, but only a few specific genes have been associated with delay discounting. Here, we aimed to identify novel genetic loci associated with delay discounting through a genome-wide association study (GWAS) using Heterogenous Stock rats, a genetically diverse outbred population derived from eight inbred founder strains. We assessed delay discounting in 650 male and female rats using an adjusting amount procedure in which rats chose between smaller immediate sucrose rewards or a larger reward at variable delays. Preference switch points were calculated for each rat and both exponential and hyperbolic functions were fitted to these indifference points. Area under the curve (AUC) and the discounting parameter *k* of both functions were used as delay discounting measures. GWAS for AUC, exponential *k*, and indifference points for a short delay identified significant loci on chromosomes 20 and 14. The gene *Slc35f1,* which encodes a member of the solute carrier family of nucleoside sugar transporters, was the only gene within the chromosome 20 locus. That locus also contained an eQTL for *Slc35f1*, suggesting that heritable differences in the expression of that gene might be responsible for the association with behavior. The gene *Adgrl3*, which encodes a member of the latrophilin family of G-protein coupled receptors, was the only gene within the chromosome 14 locus. These findings implicate novel genes in delay discounting and highlight the need for further exploration.

## Introduction

Delay discounting is the neurobehavioral process by which individuals devalue delayed rewards. It is usually assessed by measuring relative preferences between smaller rewards available immediately and larger rewards with delayed delivery.^1^ It has been equated with impulsivity, a multifaceted construct represented by several behavioral phenotypes and linked to substance use disorders (SUD).^2,3^ Recent work has called into question the utility of impulsivity as a unitary construct due to its multifaceted operational definitions.^4–7^ However, researchers in the field do not dispute the importance of the delay discounting phenotype in SUDs, only whether greater discounting should be interpreted as indicating “impulsiveness.” The lack of controversy regarding the role of delay discounting in SUDs is attributed to the extensive body of evidence accumulated over the last 25 years, with over a hundred published studies comparing delay discounting in drug users and nonusers. Over 80% of these studies have reported higher levels of delay discounting in individuals meeting criteria for SUD, and not a single published study has shown the opposite relationship.^8,9^ Higher levels of discounting have also been associated with other psychopathologies that often co-occur with SUDs, including depression, bipolar disorder, schizophrenia and attention deficit hyperactivity disorder.^10–12^ Positive associations have also been reported with pathological gambling,^13–15^ and obesity,^16,17^ suggesting a broader relationship between heightened delay discounting and psychopathology. Indeed, the pervasive association between heightened levels of delay discounting and psychopathology has led some to characterize delay discounting as a “transdisease” or “transdiagnostic” marker and high levels of delay discounting as indicative of a causal “reinforcer pathology.”^18–20^

While the link between delay discounting and substance use is well-established, the processes underlying this relationship have not yet been fully elucidated. One contributory mechanism may be common genetic substrates. Familial and twin studies, as well as genome-wide association studies (GWAS), have established that there is a genetic component to delay discounting. Twin studies have shown higher correlations within monozygotic twins compared to dizygotic twins, indicating a strong genetic contribution to the trait.^21,22^ Furthermore, in the largest human GWAS of delay discounting to date, which included 23,127 participants of European ancestry, genotype accounted for 12% of the variance of delay discounting, as measured by the Monetary Choice Questionnaire.^23^ The heritability of delay discounting in rodents has also been demonstrated using panels of inbred strains.^24–26^

These studies indicate a genetic component to discounting, but only a single gene (*GPM6B*) has ever show a genome-wide significant association with delay discounting.^23^ Other studies have identified risk genes for impulsivity as the broadly defined construct,^7,27,28^ but these genes did not have associations with delay discounting in Sanchez-Roige *et al.* (2018). This lack of concordance underscores the modest overlap between questionnaire measures of impulsivity and delay discounting, and the broader uncertainty over the true relationship between delay discounting and complex neurobehavioral traits.^4,5,29^ Several studies using animal models have examined the effects of single gene mutants, but with mixed success. No differences were reported between knockouts and wildtypes for *Lphn3*^30^ or D_4_ receptor deficiency;^31^ though reduced delay discounting was reported for conditional knockouts of *Ant1*^32^ and augmented discounting was reported following viral vector knockdown of D_2_R localized in the ventral tegmental area.^33^

Identifying the genes associated with delay discounting may provide valuable information about the transdiagnostic links between discounting and SUDs, or even psychopathologies more generally. Gene identification may suggest novel intervention targets, as well as novel indicators of heightened risk for dysregulated behavior. Furthermore, gene identification may point to critical cell types and neurocircuits that mediate differences in delay discounting and the correlated psychopathologies. Accordingly, the current study aimed to identify genes associated with delay discounting. To accomplish this aim, we phenotyped rats from a Heterogeneous Stock (HS) population. HS rats are an outbred population derived from eight inbred founder strains and have been used extensively for GWAS of other phenotypes.^34–37^ The high level of both genetic and phenotypic diversity of these rats makes this an ideal population to investigate complex neurobehavioral traits such as delay discounting and to identify associated genetic variants.^38,39^

There is some debate about the most appropriate way to quantify delay discounting.^40–42^ Changes in relative preference for the smaller, sooner versus the larger, later rewards are typically examined over a series of delays. Traditionally, functions are fitted to the points at which preferences shift at each delay (indifference points) and the slope of this function is used as a measure of delay discounting. Steeper slopes indicate heightened discounting. The function fitted most often is a hyperbolic function from early work by Mazur (1987) but other functions have also been examined.^43–46^ Furthermore, to circumvent discussions about which function is the most appropriate, others have calculated the area under the discounting "curve,” as derived from the indifference points, because this approach is function-free.^47–49^ In acknowledgement of the ongoing discussions about the best metrics to quantify delay discounting, and to take advantage of our relatively large dataset, we adopted three quantification methods in our study: the area under the discounting curve (“AUC”), the slope of the hyperbolic function (“hyperbolic *k*”), and the slope of an exponential function (“exponential *k*”) fitted to the indifference points. The exponential function was included because Mitchell *et al.* (2023) suggest that it can describe data for individual rats almost as well as the hyperbolic function based on corrected values of the Akaike Information Criterion (AIC).^50–53^ These results, combined with the focus on exponential functions from economists,^54–56^ led to the inclusion of the exponential function in our study to identify genes associated with delay discounting. Additionally, functions were fitted with and without a bias term, which captures the side preference of individual rats during the experimental trials. Side bias is not included in assessments of delay discounting used with human participants but improves the fit of the hyperbolic and exponential functions in rodent studies. The underlying drivers of bias may be multifactorial^57^ and including a side bias constant does alter the derived hyperbolic and exponential slopes.

## Methods

### Animals

Subjects were male and female genetically heterogeneous stock (HS) rats (official designation: NMcwiWFsm:HS #13673907, RRID:RGD_13673907). HS rats were purchased from Wake Forest University and arrived at Oregon Health & Science University (OHSU) in six shipments between October 2018 and February 2020. Rats from the first four shipments, cohorts 1-4, were phenotyped (N = 395). Due to the pandemic lockdown, rats from the fifth and sixth shipments, cohorts 5 and 6, could not be phenotyped but were used as breeders to generate more rats. Breeding took place at OHSU in May 2020 due to the pandemic, with instruction from Dr. Solberg Woods to preserve the genetic diversity and ensure phenotyping occurred with similarly ages rats. The offspring from this breeding were labeled as cohort 7. In total, cohorts 1-4 (N = 395) and 7 (N = 255) were phenotyped for delay discounting. These cohort groups were used as covariates in the genetic analysis below.

Rats of the same sex were pair-housed with lights on from 6:00 to 18:00 hours, at a temperature of 70°F with ad libitum access to water (except as specified below). They were transported to the laboratory for behavioral testing 5-7 days/week in squads that remained in the laboratory for approximately 2 hours. Testing occurred between 9:00 and 17:00 h, i.e., during the light phase, and rats were water restricted while in the laboratory. Rats were food restricted starting 1 week prior to the beginning of behavioral training and maintained at approximately 90% ad libitum weight by supplemental feeding immediately after behavioral sessions (PicoLab® Laboratory Rodent Diet 5L0D pellets). Weights were monitored daily before behavioral sessions. Rats were treated in compliance with the Guide for the Care and Use of Laboratory Animals^58^ and the experimental protocols were approved by the Institutional Animal Care and Use Committee at OHSU (IACUC; IP00001663).

### Apparatus

The operant chambers used to examine delay discounting were configured in a similar way to those described previously.^59^ Briefly, the modular rat operant chambers were housed in sound-attenuating chambers (Med Associates Inc., St Albans VT, USA). On one wall of the chamber there were two nonretractable levers, with a stimulus light above and a liquid receptacle below each. Between the levers was a nose poke with a light. On the opposite wall there was a speaker-tone generator combination and a clicker. Two 3.33 rpm syringe pumps were used to deliver 10% w/v sucrose solution to each of the liquid receptacles inside the chamber. MED-PC V software (1 ms resolution) was used to control the equipment and record activity. Operation of the equipment was tested prior to sessions.

### Delay Discounting Assessment

After training, rats were exposed to the adjusting amount procedure (**Figure 1A**), as adapted from Richards *et al.* (1997), and used extensively with rats.^25,57,60–63^ Briefly, sessions included free- and forced-choice trials, and ended after 60 free-choice trials occurred, or 60 minutes had elapsed. On free-choice trials, the size of the reinforcer delivered when the delay lever was pressed was 150 µl, while the reinforcer associated with the immediate lever was adjusted throughout the experimental session (initial size: 75 μl). Choice of the delay lever increased the current size of the immediate reinforcer by 10% on the following trial to a maximum 300 µl. Choice of the immediate lever decreased the current size of the immediate reinforcer by 10% on the following trial to a minimum of 5 µl. The size of the immediate reinforcer was not altered following forced-choice trials, which occurred after two consecutive choices of either the delay or immediate lever. Variable length timeouts between trials ensured that trials occurred every 30 s, regardless of the choice on the previous trial. Choice of the delay lever resulted in reinforcer delivery after 0, 2, 4, 8, 16, or 24 s. This delay remained constant within a session but varied between sessions according to a Latin Square design that was the same for all rats. Rats experienced each delay on 6 occasions.

**Figure 1.**
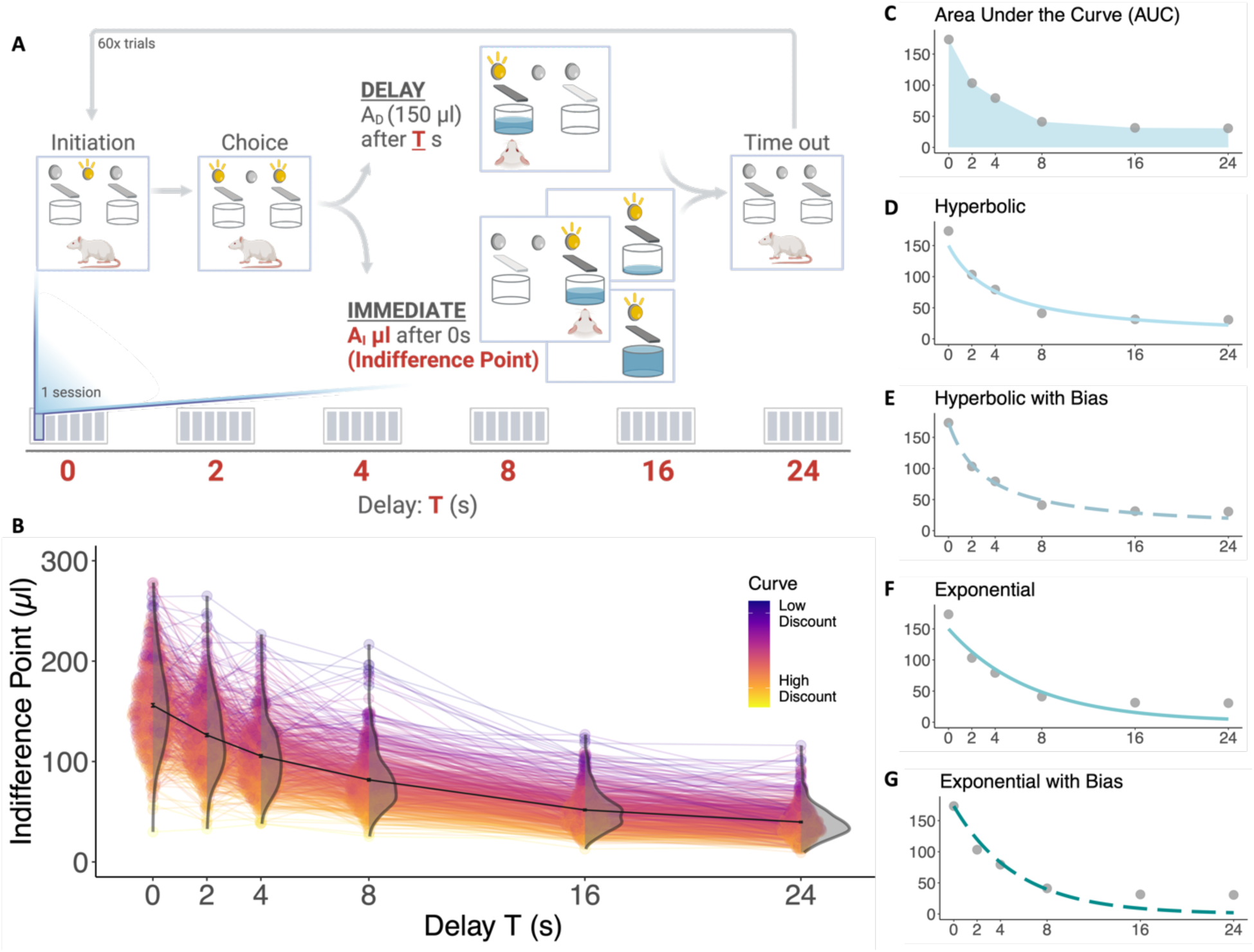
Multiple indices of delay discounting were calculated for each HS rat (n = 629). A) Schematic of the adjusting amount procedure, where one session for a specific delay includes ∼60 trials, and rats are exposed to six sessions per delay. Indifference points for each delay (T seconds) were calculated based on the A_I_ values from the last 30 trials of each session. B) Indifference points were plotted for each delay to create discounting curves for each individual rat (one line per rat). Low to high discounting, based on the steepness of the discounting curve for a rat, is denoted by color, with a low discounting rat having a darker purple curve and a high discounting rat having a brighter yellow curve. The black curve connects the mean values for each delay showing the average delay discounting curve for all rats. Error bars represent standard error. Violin plots show the distribution of indifference points for rats at each delay. C-G) Examples for delay indices are represented with a curve for a single rat. C) Area under the curve (AUC) was calculated for each rat by summing the area of the trapezoids created by the indifference points. D-G) Hyperbolic and exponential functions with and without bias were fitted to the curve for each rat. Inclusion of the bias term improved function fits, while functions without bias equalized the 0-s starting point across all rats at 150 µl.

### Statistical analysis for phenotyping

For each subject, an “indifference point” was calculated using methods described in detail in Mitchell *et al.* (2023).^53^ We used two procedures to enhance the robustness of our indifference point measures. First, we excluded any session on which fewer than 45 of the 60 free-choice trials were completed by a rat: 1,714 out of 23,400 (7.3%) sessions (650 rats x 6 delays x 6 occasions). Second, we excluded any completed session on which choices during the second half of the session were primarily on one lever (operationalized as 80% or more of trials 31-60 during the session): 967 out of the 21,686 (4.5%) completed sessions. These exclusions resulted in 20,719 sessions of data from which indifference points were derived. These indifference points provide an index of the subjective value of the 150 µl 10% sucrose solution and are expected to decrease as the delay to receiving the 150 µl reward increases, reflecting the putative decrease in its subjective value.

Two strategies were used to quantify the extent of decreases in subjective value. First, we fit either a hyperbolic^43^ or an exponential^64^ mathematical function to the indifference points for individual rats:

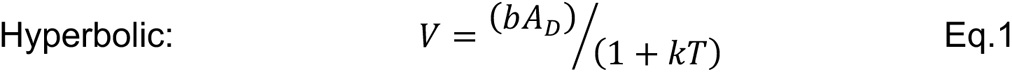

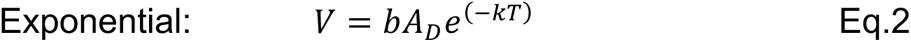

*V* represents the subjective value at the indifference points, *b* represents an individual’s side preference in the apparatus, the *A_D_* represents the amount of the larger reward (150 µl), *T* represents the time delay to that reward, *k* represents the discounting parameter (slope of the function). Importantly, side preference, or bias, *b* is calculated by dividing the indifference point at delay 0 s by 150 µl, to generate a unit free constant. Bias accounts for any preference for the left or right lever, and preferences for each were roughly equal across rats. The inclusion of bias affects the fit of both functions. The models that account for bias are better fit than assuming no bias, but the numerical value of the slopes of both functions are altered and the genetic basis for side bias remains unclear. Accordingly, both functions were fit with and without bias to calculate the *k* values (here, we designate the difference in *k* values using *“k_bias_”* and *“k_w/obias_”*). Second, we calculated the area under the discounting curve (AUC) by summing the areas of the trapezoids created by indifference points.^48^ Taken together, AUC, exponential and hyperbolic *k_bias_* and *k_w/obias_,* as well as the indifference points at each delay were all used to index the delay discounting trait for GWAS. Analysis was done in R and can be reproduced using the pipeline available on Github (https://github.com/Palmer-Lab-UCSD/HSrat_delaydiscounting).

### Genotyping

A total of 650 experimental HS rats were genotyped. Spleens were collected postmortem and used as a source of DNA for genotyping (dx.doi.org/10.17504/protocols.io.6qpvr665ovmk/v1). Spleen tissue samples were cut and processed (dx.doi.org/10.17504/protocols.io.36wgq7nryvk5/v1), and DNA was isolated using the Beckman Coulter DNAdvance Kit at the University of California San Diego (dx.doi.org/10.17504/protocols.io.8epv59reng1b/v1). All samples were normalized and processed in a randomized order prior to library preparation (dx.doi.org/10.17504/protocols.io.261genw5dg47/v1), and multiplexed sequencing libraries were prepared using the iGenomX RipTide kit (dx.doi.org/10.17504/protocols.io.j8nlkkm85l5r/v1). Final QC was performed on sequencing libraries; sequencing was performed using an Illumina NovaSeq 6000 (dx.doi.org/10.17504/protocols.io.yxmvmnw29g3p/v1). Reads were demultiplexed using fgbio v1.3.0 (http://fulcrumgenomics.github.io/fgbio/) before trimming adapters using BBDuk v38.94 (https://sourceforge.net/projects/bbmap/) and quality trimming using Cutadapt v4.1.^65^ Reads were aligned to the rat reference genome mRatBN7.2 from the Rat Genome Sequencing Consortium (GCA_015227675.2 GCF_015227675.2) using BWA-mem v0.7.17.^66^

Mapped sequences were then used to construct haplotypes and impute biallelic SNP genotypes using STITCH v1.6.6.^67^ From this set of 10,684,883 SNPs, we removed all SNPs with low imputation quality scores produced by STITCH (INFO < 0.9; 2,609,890 SNPs removed). We additionally removed all SNPs with high missing rates (missing rate > 0.1; 21,900 removed), low minor allele frequencies (MAF < 0.005; 2,600,296 removed), and extreme deviations from Hardy-Weinberg Equilibrium (HWE p < 1e-10; 2,370 removed). This filtered set of 5,451,257 SNP genotypes was used for all downstream analyses.

### Phenotype data and genetic analysis

While the AUC data were relatively normally distributed (skew = 0.80, kurtosis = 1.30), the exponential and hyperbolic values were not (exponential *k_bias_* and *k_w/obias_* skew:1.73 and 3.18, kurtosis: 7.47 and 20.33; hyperbolic *k_bias_* and *k_w/obias_* skew: 1.60 and 3.10; kurtosis: 6.32 and 20.19). To address these deviations for normality, all delay discounting traits were quantile normalized separately for males and females prior to GWAS.

To examine the effects of covariates (sex, cohorts, coat color, cage, and age), we fit linear models that predicted the phenotype based on each distinct covariate. We used linear regression to remove the effects of covariates that explained more than 2% of the variance of the trait. For AUC, and exponential and hyperbolic *k_w/obias_*, the four separate shipments of rats (cohort 1-4) as well as the final group of rats bred at OHSU (cohort 7), explained more than 2% of the variance of these indices, so they were regressed out. Additionally, cohort 7 for hyperbolic *k_bias_*, and cohort 1, 2, and 7 for exponential *k_bias_* were regressed out.

Phenotypic correlations between the five measures of delay discounting (AUC, exponential *k_bias_*, exponential *k_w/obias_*, hyperbolic *k_bias_,* and hyperbolic *k_w/obias_*) were determined using Spearman’s test. These correlations were visualized using Seaborn Clustermap, which also computes average linkage hierarchical clustering for the traits. We also included the indifference points for each delay and the bias term in the phenotypic correlation and clustering because these measures were also included in the genetic analysis.

GWAS analysis was performed using the mixed linear model analysis (MLMA) function from the Genome-wide Complex Trait Analysis (GCTA) software package to compute association statistics explaining the genetic contribution to phenotypic variance.^68^ This algorithm builds a genetic relationship matrix (GRM) between individuals and uses the leave one chromosome out (LOCO) method, which leaves out SNPs on the same chromosome as the test SNP to avoid proximal contamination.^69^ SNP heritability estimates were obtained using the restricted maximum likelihood (REML) approach from the GCTA package, which relies on the GRM to estimate the proportion of phenotypic variance explained by all SNPs.^68^ Genetic correlations between traits were estimated using bivariate GCTA-REML analysis. These correlations were also visualized and clustered using hierarchical clustering.

Genome-wide significance thresholds (α = 0.05 and 0.10) were calculated using permutation tests.^35,70^ The same thresholds were used for all delay discounting indices because all phenotypes were quantile normalized and thus had identical genotypes and identical phenotypic distributions. We report all SNPs with p-values exceeding the significance threshold of –log10(p) = 5.58 (α = 0.05) or –log10(p) = 5.36 (α = 0.10).

GWAS was only performed for rats for which all delay discounting indices could be obtained (n = 629, females = 319, males = 310), including AUC, and exponential and hyperbolic *k_w/obias_ and k_bias_*. Additional analyses were also conducted for the indifference points at each delay (2, 4, 8, 16, 24 s), and the bias term, which is based on the indifference point at the 0-s delay. Each chromosome was scanned to identify quantitative trait loci (QTLs) containing at least one SNP that exceeded the threshold. Linkage disequilibrium (LD) intervals for each significant QTL were determined by finding additional significant SNPs within 0.5 Mb that had a high correlation (r^2^ = 0.6) with the peak SNP. We generated a porcupine plot combining the Manhattan plots for the traits showing QTLs of genome-wide significance, as well as Regional Association Plots for the significant QTLs for each trait.

## Results

### HS Rat Phenotyping for Delay Discounting

Adult HS rats completed the adjusting amount procedure to measure delay discounting (**Figure 1A**) and indifference points were recorded for each time delay. Due to missing data from incomplete sessions, indices of delay discounting could not be calculated for 21 genotyped rats, resulting in a total of 629 rats (females = 319, males = 310) with complete delay discounting profiles. Indifference points were plotted to generate discounting curves for each rat **(Figure 1B**). Discounting curves for rats ranging from low to high discounting are denoted by color. In addition to indifference points at each delay, bias was calculated using the indifference point at 0-s delay as a measure of side preference. For each rat discounting curve, several indices were calculated as measures of delay discounting, including area under the curve (AUC, **Figure 1C**). Hyperbolic and exponential functions were fit to indifference points, first without bias (**Figure 1D, 1F**) and then including the bias term in the function (**Figure 1E, 1G**). Accounting for bias improved the fit of both the hyperbolic and exponential functions, while the functions without bias equalized the 0-s starting point for all rats at 150 µl.

The discounting parameter, *k*, was calculated for the exponential and hyperbolic functions with and without bias for each rat. These five values (AUC, exponential *k_bias_*, exponential *k_w/obias_*, hyperbolic *k_bias_,* and hyperbolic *k_w/obias_*) were used as the measures of delay discounting in GWAS. We subsequently included the indifference points at each delay as well as the bias term in the downstream genetic analysis to determine if the individual components making up the delay curve were driving the results. Strong phenotypic correlations were found between the exponential and hyperbolic *k* parameters, while AUC had stronger phenotypic correlations with the indifference points and clustered separately (**Figure 2A**). Though there were 12 phenotypic indices total, two phenotypic clusters emerged due to strong correlations between the measures. Genetic correlations followed a similar pattern of clustering (**Figure 2B**). SNP heritability estimates ranged from 0.08 ± 0.05 (bias and delay 0) to 0.19 ± 0.06 (AUC and exponential *k_w/obias_*; **Figure 2B**). The three delay discounting indices without bias (AUC, exponential *k_w/obias_*, and hyperbolic *k_w/obias_*) had higher heritability estimates than the two delay indices with bias (**Figure 2B, Supplemental Table 1**). Heritability estimates for AUC, exponential *k_w/obias_*, and hyperbolic *k_w/obias_* were consistent with many other behavioral traits that have been reported previously in HS rats in the past. However, estimates for the indifference points at each delay and the bias term were generally lower than the composite delay discounting indices.

**Figure 2.**
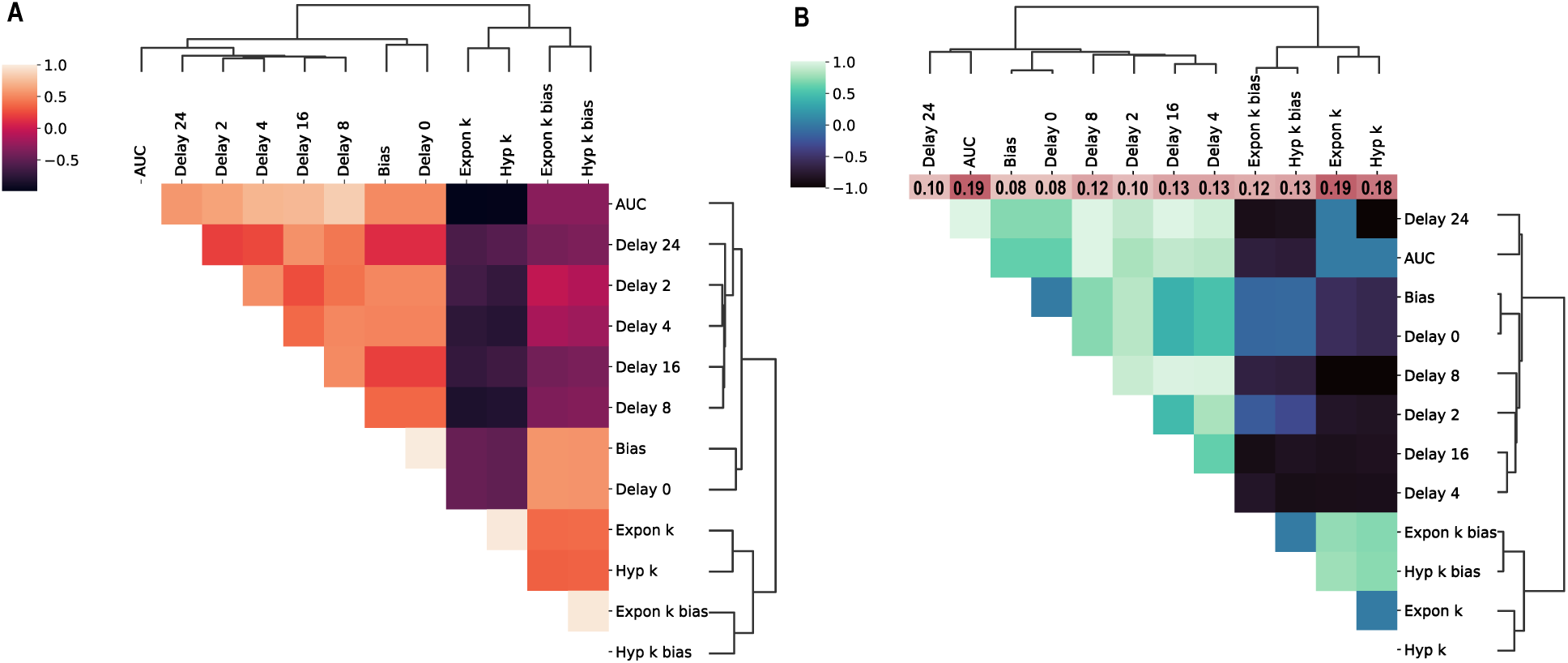
Phenotypic and genetic correlations were calculated for all delay discounting measures. A) Phenotypic correlations were determined using Spearman’s test, and traits were clustered using average linkage hierarchical clustering. B) Genetic correlations were calculated using bivariate REML analysis implemented in GCTA. SNP heritability estimates for each trait are denoted in the color bar above the heatmap.

### GWAS for Delay Discounting

After filtering and controlling for quality as described in the Methods section, we obtained genotypes at 5,451,257 SNPs for 629 rats. We performed a GWAS of delay discounting indices for AUC, exponential *k* and hyperbolic *k* parameters with (*k_bias_*) and without bias (*k_w/obias_*) for 629 rats. Indifference points for each delay and the bias term were subsequently included in the GWAS.

We detected a genome-wide significant locus on chromosome 20 for both the AUC and exponential *k_w/obias_* delay curve indices **(Figure 3A-C),** but not for hyperbolic *k_w/obias_*, though this same locus was almost significant (-log_10_(p) = 4.915). GWAS identified the same top SNP (20:32,221,020) in the locus for AUC and exponential *k_w/obias_*, which had a minor allele frequency of ∼17%. The minor allele was derived from BN/N and ACI/N, whereas the other 6 founders had the major allele. The minor allele was associated with higher discounting, as demonstrated by lower AUC (**Figure 3E**) and higher values of exponential *k_w/obias_* **(Figure 3F**). For AUC, the top SNP showed a -log_10_(p) = 5.689, which corresponds to a *p*<0.05. For exponential *k_w/obias_,* the same SNP showed a -log_10_(p) = 5.449, which corresponds to p<0.10. For hyperbolic *k_w/obias_,* this SNP showed a -log_10_(p) = 4.915, which was near threshold but not significant. A porcupine plot combining the two traits on is shown in **Figure 3A**. This top SNP is located in an intron of the gene *Scl35f1* as depicted in the locus zoom plots in **Figure 3B and 3C**, and the nearby SNPs in this QTL that are in strong linkage disequilibrium (LD) with the top SNP (r^2^ > 0.8) are located in multiple exons and introns of *Slc35f1*. *Scl35f1* encodes a member of the solute carrier family 35, which has been implicated in brain-related function and neurodevelopmental disorders. There are two expression QTLs (eQTL) for *Slc35f1* (20:32,306,446 and 20:32,306,658) that reflect heritable differences in expression of *Slc35f1* in whole brain and prelimbic cortex. These eQTLs are in strong LD with the top SNP (r^2^ = 0.97), suggesting that heritable differences in the expression of *Slc35f1* may mediate the effect of this locus on delay discounting. We did not identify any other eQTLs in this locus nor were they any coding variants that were predicted to have major effects of in this locus. There were no significant loci associated with exponential or hyperbolic indices with bias (*k_bias_*).

**Figure 3.**
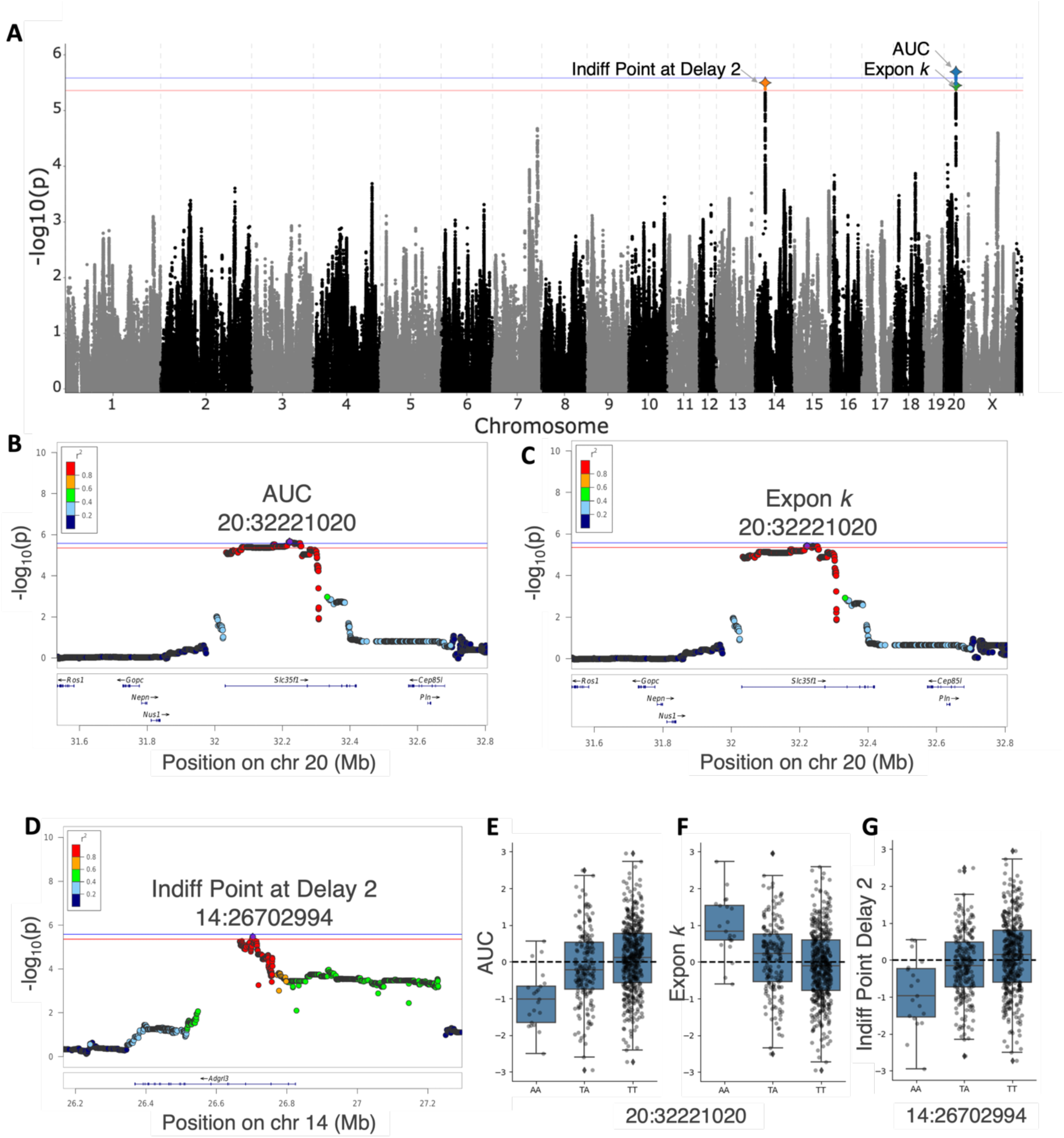
GWAS for AUC and exponential *k_w/obias_* measures for delay discounting, as well as the indifference point at the 2 s delay resulted in significantly associated loci mapping to the two genes: *Slc35f1* and *Adgrl3*. A) The porcupine plot displaying the chromosomal distribution of all p-values combines the three delay discounting measures and shows both significant chr 20 SNPs and the chr 14 SNP. The blue and red lines show the significance thresholds derived from the permutation tests: -log_10_(p) = 5.58 (alpha = 0.05) and -log_10_(p) = 5.36 (alpha = 0.10), respectively. B-D) Regional association plot for the QTL on chr 20 for both delay discounting indices: AUC and exponential *k_w/obias_*, and for the QTL on chr 14 for indifferent point at 2s delay. The x-axis depicts chromosomal position in Mb and the y-axis shows the significance of the association (-log_10_(p)), and individual dots represent SNPs. Purple denotes the top SNP associated with the trait, and color of the dots indicate level of linkage disequilibrium between top SNP and other nearby SNPs. E-G) Effect plots for all three measures showing the genetic effect of the peak SNP. The minor allele for the chr 20 SNP was associated with heightened discounting with lower AUC and higher exponential *k* values, and the minor allele for the chr 14 SNP was associated with lower indifference points at 2 s delay.

GWAS also detected a genome-wide significant locus on chromosome 14 for the indifference point at the 2 s delay (**Figure 3A**). The SNP 14:26,702,994 (-log_10_(p) = 5.496, p<0.10) had a minor allele frequency of ∼21% and the major allele was derived from MR/N, with all other founders having the minor allele. Lower indifference points for the short delay were associated with the minor allele (**Figure 3G**). Of note, this SNP neared genome-wide significance for both the bias term and indifference point at 0s delay (-log_10_(p) = 4.418), which are perfectly correlated with each other as bias is calculated from this indifference point. This SNP is located in an intron of the gene *Adgrl3*, which encodes a type of G-protein coupled receptor. Additionally, multiple eQTLs exist in the chromosome 14 locus for *Adgrl3* for various tissues including brain (14:26,678,469), infralimbic cortex (14:26,745,338), and lateral habenula (14:26,724,419), which all were in high LD with the top SNP (r^2^ > 0.96). Multiple other eQTLs for *Adgrl3* exist in this locus that are also in LD with the top SNP, but to a lesser degree (r^2^ ∼ 0.6), and these include: basolateral amygdala (14:26,776,241), nucleus accumbens (14:26,782,299, 14:26,779,935), orbitofrontal cortex (14:26,776,241), and prelimbic cortex (14:26,790,378, 14:26,776,241).^71^

## Discussion

The main aim of the present study was to identify genes associated with delay discounting using genetically heterogeneous rats. To accommodate the lack of consensus about discounting curve models, delay discounting was indexed in multiple ways including fitting a hyperbolic and an exponential function to the indifference points both with and without a bias term, as well as calculating the area under the indifference points. GWAS for these indices of delay discounting, as well as the bias term and indifference points at each delay, identified two genome-wide significant QTLs located on chromosome 14 and 20, despite the relatively small sample size. The gene *Scl35f1*, which encodes a member of the solute carrier family of membrane transport proteins,^72^ was the only gene within the chromosome 20 locus; and the *Adgrl3* gene, which encodes a member of the latrophilin family of G-protein coupled receptors,^73^ was the only gene within the chromosome 14 locus. There were also multiple eQTLs for tissues in the brain for *Slc35f1* and *Adgrl3* that were in high LD with the top SNPs for these loci (r^2^ > 0.96),^71^ meaning that these loci also change expression of *Slc35f1* and *Adgrl3*, which may be driving the observed behavioral differences.

SNP heritabilities, or the proportion of variance accounted for by SNPs, for the delay discounting indices AUC, exponential *k_w/obias_*, and hyperbolic *k_w/obias_*, were estimated to be 19%, 19%, and 18%, respectively. While estimates were substantially lower for exponential and hyperbolic *k* with bias (12% and 13%, respectively), as well as the bias term itself (8%). The difference in heritability among the discounting indices with and without bias is of note, however, as the inclusion of the bias term in both functions increases variance accounted for, but simultaneously reduces heritability estimates. The underlying genetics of side bias remain opaque, though, and appear to introduce noise making the genetic bases of delay discounting more difficult to distinguish.

As expected for SNP heritabilities, these heritabilities were substantially lower than estimates derived from inbred strains. For example, Wilhelm and Mitchell (2009) reported 40% heritability for hyperbolic *k* with bias in six inbred strains of male rats using the adjusting task described here (measures without bias were not reported), and Richards *et al.* (2013) reported 50% heritability for AUC in eight inbred strains of male rats using a nonstandard delay discounting task. Lower estimates have been reported in mouse studies. Isles *et al.* (2004) estimated heritability to be 16% for a small immediate versus larger later choice preference measure based on four inbred strains of male mice.^74^ In another more recent screen of heritable variation in delay discounting, Bailey *et al.* (2021) used male and female mice from the highly genetically diverse Collaborative Cross (CC) recombinant inbred panel of mice as well as the eight founder strains from which all CC mice are derived.^24^ The combined 18 strains demonstrated significant heritability for a proxy measure of delay discounting with 25% of the variance explained by strain differences.

### Slc35f1

GWAS for AUC and exponential *k_w/obias_* identified the same significant top SNP that mapped to the gene *Slc35f1*, which encodes a member of the solute carrier (SLC) family of membrane transport proteins.^72^ The SLC35 family of nucleoside-sugar transporters were thought to localize in the Golgi apparatus and endoplasmic reticulum (ER);^75^ however, a more recent study of SLC35F1 protein expression in the adult mouse forebrain did not find co-localization of *Slc35f1* with the Golgi apparatus or ER.^76^ Nevertheless, *Slc35f1* did have high expression in the soma and dendrites of neurons in numerous cortical and diencephalic structures including the hippocampus and thalamus.^76^ The authors suggested the possible involvement of *Slc35f1* in the formation and function of dendritic spines or synaptic plasticity. High *SLC35F1* mRNA expression has also been found in both fetal and adult human brain tissues.^77^ Consistent with the protein expression patterns in the murine study, the SLC35F1 protein as well as RNA is highly expressed in multiple brain regions in human tissue including the cerebral cortex, hippocampus, and amygdala (proteinatlas.org).^78,79^ The high neuronal expression and potential role in dendritic spine dynamics point to a brain-related function, yet the full molecular mechanisms by which SLC35F1 participates in neuropsychiatric behaviors and substance use disorders (SUDs) remains unresolved.

In humans, there is some evidence to suggest that *SLC35F1* is involved in critical brain pathways underlying behavior and/or neuropathophysiology, which may give some clues about its association to delay discounting. First, Szafranski *et al.* (2015) described six unrelated pediatric epilepsy patients with microdeletions within a ∼5Mb region on 6q22.1q22.23.^80^ They narrowed the critical region to a segment that included a putative cis-regulatory sequence of the *SLC35F1* promoter and a portion of the *SLC35F1* gene itself, among other regulatory and gene sequences. Importantly, patients with microdeletions spanning this *SLC35F1* regulatory region had a constellation of varying presentations including multiple types of recurrent or refractory epilepsy, autism spectrum disorder, speech and language delay, abnormal EEG, cognitive delay, developmental regression, intellectual disability, tremor, and global delay. Additionally, Fede *et al.* (2021) recently described a patient with an *SLC35F1* gene mutation who exhibited a Rett syndrome (RTT)-like phenotype where she experienced refractory seizures, had severe intellectual disability and limited speech, and was unable to walk independently.^81^ Together, these genetic case studies suggest an important neurodevelopmental role for *SLC35F1*.

Several human GWAS for neurological and psychiatric phenotypes have also implicated *SLC35F1.* In a GWAS meta-analysis of attention-deficit/hyperactivity disorder (ADHD) and bipolar disorder, SNPs annotated to *SLC35F1* neared significance, but were subthreshold (best *p* = 6 x 10^-8^).^82^ Interestingly, however, the strongest SNP identified in this study was rs11756438, which was in LD with SNPs in the *SLC35F1* gene. Another SNP located in *SLC35F1* also neared significance (*p* = 3 x 10^-6^) in a recent GWAS of schizophrenia,^83^ which is noteworthy in light of the known connection between steeper discounting and schizophrenia.^10^ Furthermore, in an updated GWAS meta-analysis of educational attainment with about three million individuals, SNPs rs11755280 and rs12213071 located in the *SLC35F1* gene were significantly associated with educational attainment (*p* = 2 x 10^-9^ and *p* =1.26 x 10^-9^, respectively).^84^ Four other SNPs mapped to genes in the SLC35 family were also significant, including *SLC35D2, SLC35F4,* and *SLC35F5*. Importantly, lifetime outcomes such as educational attainment have been negatively associated with delay discounting, where individuals with steeper delay discounting have lower educational attainment.^85^

Genetically altering *SLC35F1* in murine model systems has been less successful at recapitulating the phenotpyes observed in humans. Deletion of *Slc35f1* in mice did not result in any phenotypic outcomes related to the RTT-like syndrome described by Fede *et al.* (2021), nor the microdeletion syndrome described by Szafranski *et al.* (2015).^86^ However, those studies did not assess delay discounting or similar behavioral constructs.

### Adgrl3

GWAS for the indifference point at the 2 s short delay identified a genome-wide significant locus on chromosome 14 that mapped to the gene *Adgrl3*, and there were several eQTLs for *Adgrl3* at loci in strong LD with the top SNP. *Adgrl3,* also called *Lphn3*, encodes a member of the latrophilin (LPHN) subfamily of G-protein coupled receptors.^73^ The human homolog has been reliably associated with ADHD,^87,88^ and has been suggested to confer susceptibility to SUDs using a tree-based predictive analysis.^89^ Importantly, delay discounting, ADHD, and SUDs are all genetically corelated with one another and often co-occur.^23,90,91^ Functionally, *ADGRL3* is involved in synapse development in the cortex^92^ and is highly expressed in the brain (proteinatlas.org).^79^ Unfortunately, as mentioned before, in a study using a delay discounting task where rats were given an option between immediate food pellets or delayed, but a larger number of food pellets, *Lphn3* knockout rats did not show any differences in what the author’s term “impulsive choice” compared to wildtype controls.^30^ However, in a different rat study, *Lphn3* knockout in Sprague-Dawley rats did result in hyperactivity, increased acoustic startle response, and reduced activity in response to amphetamine administration.^93^ Furthermore, loss of the homologous gene (*lphn3.1*) in zebrafish resulted in ADHD-like behavior and abnormal development of dopaminergic neurons,^94^ and *Lphn3* null mutant mice showed hyperactivity and increased sensitivity to cocaine.^95^ More recently, however, *Adgrl3* knockout in mice on a B6/J background showed no neuro- or behavioral differences,^96^ though delay discounting was not assessed.

*Adgrl3* has also been associated to multiple traits in human GWAS. Several variants have been associated with educational attainment, including in a recent GWAS using data from the UK Biobank (*p =* 5 x 10^-11^).^97,98^ *Adgrl3* has also been associated with risk-taking behavior (*p* = 1 x 10^-9^),^99^ smoking initiation (*p* = 5 x 10^-9^),^100^ and externalizing behavior (*p* = 5 x 10^-9^).^101^ Considering its association to ADHD and potential SUD risk, as well as evidence in model organisms for its involvement in neuropsychiatric-related behaviors, *Adgrl3* is a strong candidate for further study.

### Limitations and conclusions

While our study yielded intriguing results that will form a foundation for future research on the genetics of delay discounting, the work is limited by its small sample size.

This may have contributed to our inability to replicate the findings of the recent human GWAS of delay discounting identified a significant association with the SNP rs6528024, which is located on the X chromosome in an intron of the neuronal membrane glycoprotein M6B gene (*GPM6B)*.^23^ In a follow-up study, *Gpm6b* deletion in C57BL/6J mice resulted in an increased preference for smaller immediate rewards compared to larger delayed rewards, reflecting higher delay discounting.^102^ A small, preliminary rat study, using CRISPR to delete *Gpm6b*, was consistent with these data (*p* = 0.18) (Mitchell, personal communication). However, it is also possible that HS rats do not have sufficient variability in *Gpm6b* expression, in which case we would not detect *Gpm6b* even if it is truly associated with delay discounting in other populations or species. Conversely, no human GWAS of delay discounting has detected a significant association with *SLC35F1*. This may be attributable to our identification of these loci being based on different indices of delay discounting (AUC and exponential *k* without bias)^103^ that provides only a measure of hyperbolic *k* (without bias as always the case in studies with human participants) or to a lack of sufficient variation in *SLC35F1* or *ADGRL3* among humans. Future work could re-examine data from the human GWAS using these alternative metrics of delay discounting. Other differences between assessments of delay discounting in human and rodent studies may also be a factor in the lack of concordance, for example, the use of hypothetical versus real rewards in human versus rodent studies, the time scale over which discounting is assessed, as discussed by numerous authors.^104,105^ These methodological differences are difficult to address but explorations of whether such moderating effects are genetically influenced could be the focus of future studies. Finally, while not identified at genome wide significant levels, we assume that additional loci can and will be identified in the future, once larger sample sizes are available.

## Author Contributions

SHM and AAP designed the study. SHM and her lab members performed the behavioral phenotyping, and SHM and MKL completed the analyses of the behavior and quantified the phenotypes of the individual subjects. SHM, MKL, and AAP wrote the manuscript. AAP and OP oversaw the genotyping. DC, KN, KC performed the genotyping. ASC, AAP, OP, TS, and BBJ developed and oversaw the genetic analysis. SZ, SAM, and BL prepared the phenotypic data for genetic analysis. LSW and AB produced the HS rats used for this study.

## Acknowledgements

We would like to thank Jordan Bromley, Katie Garland, Skylar McShane, and Deborah Sevigny-Resetco for conducting experimental sessions to phenotype the rats. William Guethlein assisted in configuring the operant chambers for the study and in writing the MED-V programs used to collect data. He created the data processing Excel macros used to summarize the session data prior to quantification of the phenotypes. Jianjun Gao and Riyan Cheng have broadly contributed to genetic data processing standards used in the project.

