## Supplemental Table 1 for "Genome-wide association study of delay discounting in Heterogenous Stock rats"

**Supplementary Table 1.** Heritabilities of phenotypic indices for delay discounting

| Trait | Heritability | -log10(p) |
|---|---|---|
| AUC | 0.185 | 3.888 |
| Indifference point delay 16 | 0.128 | 2.584 |
| Indifference point delay 2 | 0.104 | 2.026 |
| Indifference point delay 24 | 0.101 | 1.416 |
| Indifference point delay 4 | 0.133 | 2.559 |
| Indifference point delay 8 | 0.122 | 2.011 |
| Exponential *k* (with bias) | 0.116 | 1.689 |
| Exponential *k* (without bias) | 0.186 | 3.807 |
| Hyperbolic *k* (with bias) | 0.129 | 1.933 |
| Hyperbolic *k* (without bias) | 0.179 | 3.667 |
| Indifference point delay 0 | 0.076 | 1.418 |
| Bias | 0.076 | 1.418 |
